# Structural Model of the Proline-rich Domain of Huntingtin exon-1 fibrils

**DOI:** 10.1101/2020.04.17.046714

**Authors:** A. S. Falk, J. M. Bravo-Arredondo, J. Varkey, S. Pacheco, R. Langen, A. B. Siemer

## Abstract

Huntington’s disease (HD) is a heritable neurodegenerative disease that is caused by a CAG expansion in the first exon of the huntingtin gene. This expansion results in an elongated polyglutamine (polyQ) domain that increases the propensity of huntingtin exon-1 (HTTex1) to form cross-β fibrils. While the polyQ domain is important for fibril formation, the dynamic, C-terminal proline-rich domain (PRD) of HTTex1 makes up a large fraction of the fibril surface. Because potential fibril toxicity has to be mediated by interactions of the fibril surface with its cellular environment, we wanted to model the conformational space adopted by the PRD. We ran 800 ns long molecular dynamics (MD) simulations of the PRD using an explicit water model optimized for intrinsically disordered proteins. These simulations accurately predicted our previous solid-state NMR data and newly acquired EPR DEER distances, lending confidence in their accuracy. The simulations show that the PRD generally forms an imperfect polyproline II (PPII) helical conformation. The two polyproline (polyP) regions within the PRD stay in a PPII helix for most of the simulation, whereas occasional kinks in the proline rich linker region cause an overall bend in the PRD structure. The dihedral angles of the glycine at the end of the second polyP region are very variable, effectively decoupling the highly dynamic 12 C-terminal residues from the rest of the PRD.

**Statement of Significance:** HD is caused by a polyQ expansion in the exon-1 of huntingtin, which results in the formation of fibrillar huntingtin aggregates. Although the polyQ domain is the site of the disease-causing mutation, the PRD domain of HTTex1 is important for fibril toxicity and contains many epitopes of fibril-specific HTTex1 antibodies. Here, we present a structural and dynamic model of the highly dynamic PRD domain using a combination of EPR, solid-state NMR, and MD simulations. This model paves the way for studying known HTTex1 fibril specific binders and designing new ones.

## Introduction

Huntington’s disease (HD) is a fatal neurodegenerative disease for which there is no cure. HD is caused by an expansion of a polyglutamine (polyQ) encoding tract (CAG repeats) in exon 1 of the huntingtin gene (*HTTex1*) beyond 36 repeats (1). This makes HD the most common member of a class of diseases caused by polyQ expansions (2). Postmortem examination of HD patient brains shows large β-sheet rich deposits of HTTex1 protein, which can be generated by aberrant splicing (3, 4). Likewise, HTTex1 can form fibrils in vitro, and these fibrils have been shown to be toxic to cells (5, 6).

HTTex1 can be divided into 3 domains (see **Figure 1**): a 17 residue amphiphilic N-terminal (N17) domain, the polyQ tract of variable length, and a C-terminal domain that is rich in prolines (PRD). The PRD contains two 11 and 10 residue polyproline (polyP) tracts that are linked by a proline rich sequence. The polyQ tract forms the core of HTTex1 fibrils, and an elongated polyQ domain accelerates fibril formation and disease onset (7, 8). The N17 domain has been shown to play a major role in fibril formation and adopts a helical structure in fibrils (9–11). The polyQ tract and N17 domain have highly static and intermediate dynamics respectively (12–14).

**Figure 1:**
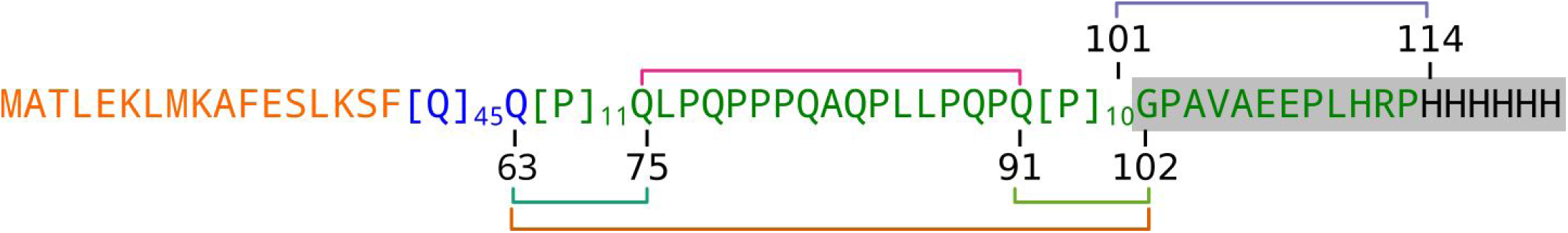
Sequence of HTTex1. The N17 region is highlighted in orange, the polyQ domain in blue, the Pro-rich C-terminus in green, and the HIS-tag in black. The 5 DEER distances measured in this study are indicated with colored brackets. The residues that were assigned site specifically using solid-state NMR are highlighted in gray.

In contrast, electron paramagnetic resonance (EPR) and nuclear magnetic resonance (NMR) studies showed that the PRD remains in a highly dynamic state even after fibril formation (12–15) and that it has essentially the same conformation in fibrils as in monomers (12). The presence of the PRD is detrimental to fibril formation by polyQ peptides (16–19) and serves as binding site for a number of proteins (20–23). It has also been shown that HTTex1 fibrils that differ in their cellular toxicity, differ in the structure and dynamics of the PRD rather than in their N17 domain or polyQ tract (5, 24, 25). Together, these findings support our previously published bottlebrush model of HTTex1 fibrils in which dynamic PRD bristles form the surface and the polyQ and N17 regions form the less accessible core (12). Fibril toxicity will always be mediated by the interaction of the fibril surface with its environment. Therefore, our goal is it to understand the structure of this fibril surface in detail. Biophysical studies and computer simulations indicated that the PRD adopts an extended polyproline II (PPII) helix (12, 13, 15, 24, 26). To test this hypothesis and to refine this model using inter-residue distances, we used a combined EPR, solid-state NMR, and molecular dynamics (MD) simulation-based approach.

The molecular description of intrinsically disordered proteins (IDPs) or intrinsically disordered domains (IDDs) is not a single structure but an ensemble of structures that describes the conformational flexibility of the protein. Both Monte Carlo and MD approaches can be used to generate such an ensemble. The quality of such an ensemble then needs to be either adjusted by selecting a suitable subset of conformers in the case of Monte Carlo simulations (27) or to be verified using experimental data in the case of MD simulations. Accurately reproducing experimental data, such as NMR relaxation rates, using MD simulations of IDPs is still challenging and a focus of active research (28, 29). One problem is that most MD force fields have been developed for globular proteins and have a tendency to produce collapsed, globular structures, typically not found in IDPs. Therefore, the choice of suitable force fields and water models is crucial for obtaining MD trajectories that are compatible with experimental data (30).

In the following, we show how double electron–electron resonance (DEER) distances can be used to select a suitable water model, force field, and starting structure for atomistic simulation of the PRD of HTTex1. Our resulting MD simulations not only correctly reproduced DEER distances but also our previously reported NMR relaxation parameters (15). The resulting conformational ensemble shows that the PRD mainly adopts a polyproline II helix, although with a high degree of flexibility and kink in the linker between the two polyproline tracts and in the residues following the second polyproline tract.

## Materials and Methods

### Protein expression and purification

HTTex1 fusion proteins were expressed, purified, and spin labeled as described previously (13, 31). In short, the thioredoxin fusion protein of HTTex1 (Trx-HTTex1) was recombinantly expressed in a pET32a vector using *E.coli* BL21 (DE3) cells. The double Cys mutants for EPR measurements were first purified using a His60 column (Clontech) followed by labeling with MTSL (1-oxyl-2,2,5,5 tetramethyl-Δ3-pyrroline-3-methylmethanethiosulfonate), and then purified using a HiTrap Q XL anion exchange column (GE Healthcare) via an AKTA FPLC system (Amersham Biosciences).

Uniformly ^13^C-^15^N labeled HTTex1 fibril samples for solid-state NMR experiments were prepared as described previously (15).

### EPR spectroscopy

Four-pulse DEER experiments (32) were done to determine the distance between spin labels. The measurements were done on a Bruker Elexsys E580 X-band pulse EPR spectrometer equipped with a 3 mm split ring (MS-3) resonator, a continuous-flow cryostat (CF935, Oxford Instruments), and a temperature controller (ITC503S, Oxford Instruments) at a temperature of 78 K. 20 µl of double spin labeled samples were adjusted to a final fusion protein concentration of 20 µM and 10% of glycerol was added as a cryoprotectant. The samples were flash frozen in liquid nitrongen before the measurement. Data were fitted using Tikhonov regularization as implemented in DEER Analysis 2019 (33).

### NMR spectroscopy

Solid-state NMR *R*_*1ρ*_ rates were measured as described previously (15). In short, a 14.1 T Agilent DD2 solid-state NMR spectrometer with a T3 1.6 mm probe was used. The MAS frequency was 12 kHz and the temperature was maintained at 0 °C. Hard pulses were done with RF-field strengths of 200 kHz and 50 kHz for ^1^H and ^15^N, respectively. *R*_*1ρ*_ relaxation dispersion was measured with 6 and 18 kHz ^15^N spin lock pulses. 2.5 kHz WALTZ ^1^H decoupling was used during detection.

### Molecular Dynamics Simulations

Simulations were run in OpenMM using the AMBER ff99SB force field along with the TIP4P-D water model. The PRD was simulated starting from an extended polyproline II helix conformation (Phi=-70.00, Psi=140.00, Omega=180.00) (34–36). The starting structure was made using the ProBuilder webserver (https://nova.disfarm.unimi.it/probuilder.htm). Assuming a 46 residue polyQ domain, residues Q63 to P113 were simulated. The N, Cα, and CO atoms of Q63 (the last glutamine residue in the HTTex1 sequence) were constrained to their starting positions by 0.5 kcal/Å restraints to simulate the impact of the PRD being attached to the static fibril core at this position. Two simulations were run for a total of 800 ns each. One with the fragment described above and another simulation with an additional 6 residue C-terminal His-tag. This was done to be consistent with both the NMR and EPR experiments that were performed with and without the HIS-tag (15). The chain was orientated along the Z-axis and centered inside a water box that was 70 x 70 x 210 Å (His-tag) or 100 x 100 x 190 Å (no His-tag). The simulations were run as an NPT (constant number, pressure, and temperature) ensemble with a temperature of 0°C and a pressure of 1 bar in order to match the conditions in which NMR experiments were performed. All proline residues were simulated starting in the trans conformation and all histidines were simulated in their uncharged form. The system was neutralized via the addition of a single sodium atom. Both simulations were run using 2 fs timesteps, with fixed hydrogen bonds, and frames were taken every 20 ps. The first 200 ns of both simulations was regarded as the equilibration time and not used in the calculation of experimental parameters. The python script used to run the simulation and an OpenMM implementation of the TIP4P-D water model, the starting structures, and all simulation outputs are available upon request.

### Calculation of experimental parameters

The program RotamerConvolveMD, which is based on the MDAnalysis python package (37–39), was used to add a set of different MTSL spin label rotamers to every 10^th^ frame of the MD trajectories and to calculate the resulting distance distribution *P*_MD_. These calculations used the MTSSL 298K 2015 rotamer library (38).

*R*_*1*_ and *R*_*2*_ relaxation rates were obtained from the simulations using the equations described by Schanda and Ernst (40). First, the correlation function for the NH bond of each residue was computed as follows (41):

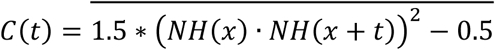

Here, *NH(x)* and *NH(x+t)* are the normalized N-H vectors at time *x* and time *x+t*. The overbar indicates that C(*t*) is averaged for all possible time points *x* during the simulation (41). Because there are fewer timepoints to average when *t* is larger, the correlation function was calculated for *t* = 0 ns to *t* = 400 ns. Afterwards the correlation function for each residue was used to fit a model free, biexponential decay function:

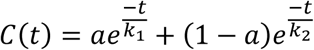

where *a* is the relative weight of the two exponential decays with rates *k*_*1*_ and *k*_*2*_. Using this model free approach and the fact that the order parameter in the dynamic PRD is *S*^*2*^≈0, the spectral density function (*J(ω*), i.e. the Fourier transform of the correlation function) can be calculated using the following equation:

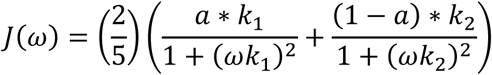

where ω is a frequency. Finally, the spectral density function was used to predict the relaxation rates *R*_*1*_ and *R*_*2*_, using the following equations:

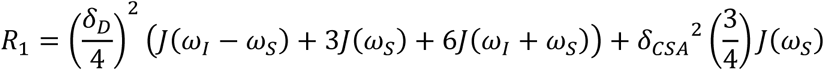

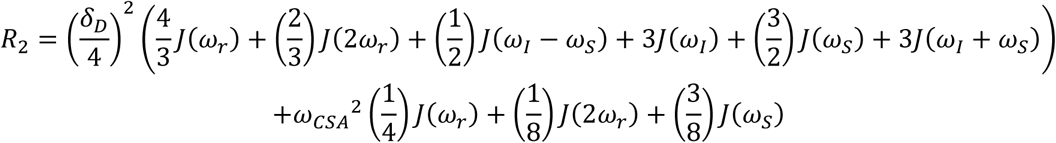

Where *δ*_D_ is the dipolar coupling between the amide proton and nitrogen, *δ*_CSA_ is the ^15^N chemical shift anistropy, *ω*_I_ is the ^1^H Larmor frequency, ω_S_ is the ^15^N Larmor frequency, and *ω*_r_ is the MAS frequency (40). The fit of the correlation function and the calculation of ^15^N *R*_*1*_ and *R*_*2*_ were done using an in-house Mathematica script that is available upon request.

Chemical shifts for each frame of the simulations after 200 ns of equilibration were calculated using the program SHIFTX2 (42) and the resulting shifts were averaged over the entire simulation. The chemical shifts were then converted into secondary chemical shifts by subtracting site-specific random coil chemical shifts calculated using the program POTENCI (43).

### Analysis of MD trajectory

The MDAnalysis (0.20) python package (37) (https://www.mdanalysis.org/) was used to calculate dihedral angles, Cα-Cα distances, and K-means clustering of the MD trajectories using in-house python scripts that are available upon request.

## Results

### Measurement of EPR distances

As a reference data set for our simulations, we determined overall distance distributions within the C-terminus of the HTTex1 fusion protein using DEER EPR. We measured 5 distances within the C-terminus as indicated in **Figure 1**: between 63R1 and 75R1 (where R1 refers to the spin-labeled side chain) in the first polyP stretch (P11); between 75R1 and 91R1 in the proline-rich linker between the two polyP stretches (L17); between 91R1 and 102R1 in the second polyP stretch (P10); between 101R1 and 114R1 in the C-terminal sequence (C12); and between 63R1 and 102R1, as a measure of the extension of the PRD. The DEER data, resulting distance distributions *P*_DEER_, and the mode (i.e. the most frequent distance) of these distributions are shown in **Figure 2**.

**Figure 2:**
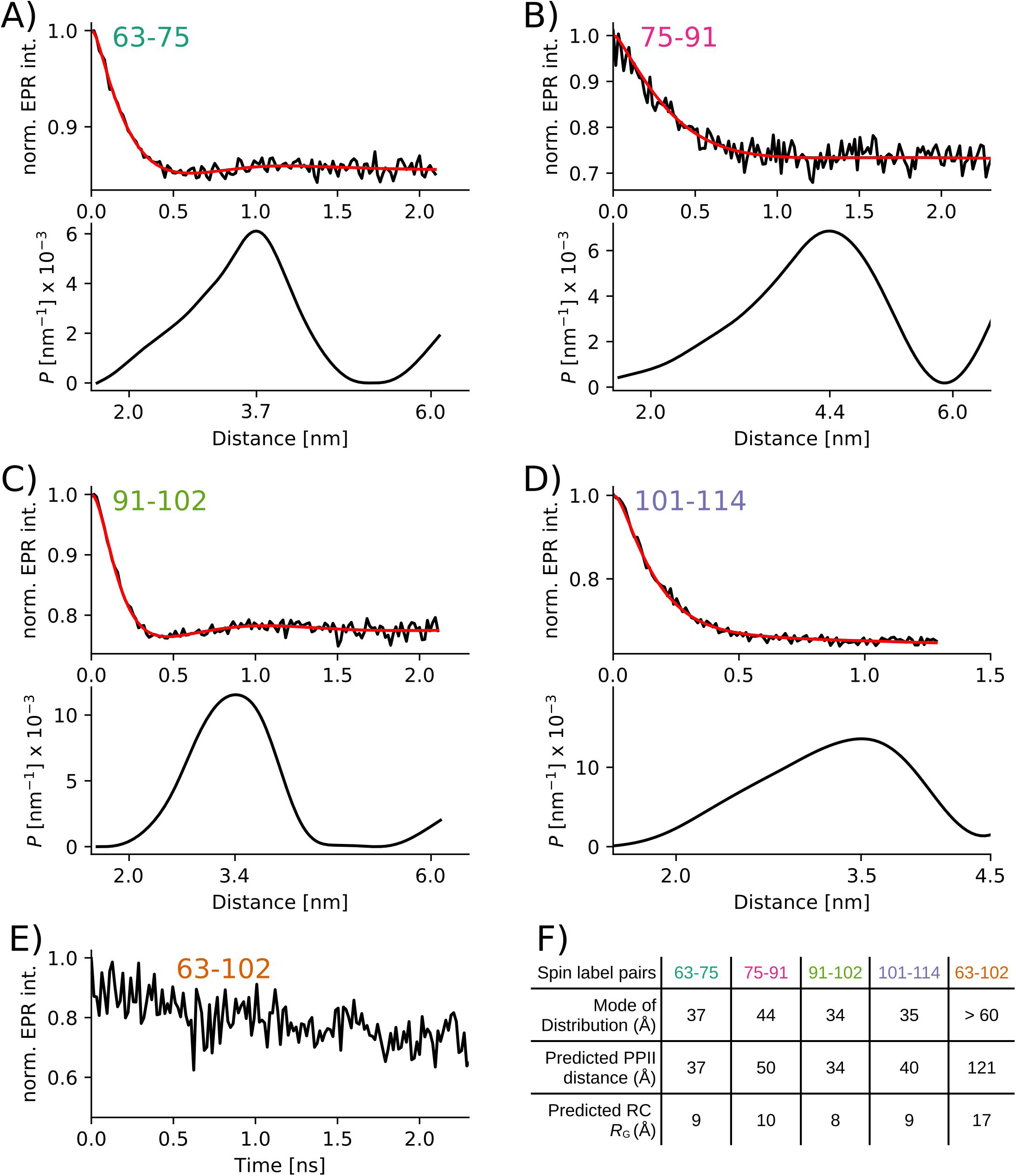
Distance distributions in the C-terminus of HTTex1 fusion protein measured via EPR DEER spectroscopy. A-D) Top panels: baseline corrected DEER data (black) and fit using Tikhonov regularization (red). Bottom panels: distance distribution P_DEER_ corresponding to fit. The mode of the distribution is indicated using a tick mark. E) Same as A-D but because the distance between 63-102 was above the detection limit, only the raw data without baseline subtraction and fit is shown. F) Table listing mode of individual distances distributions. For comparison, the theoretical distances for a polyP II helix assuming an increase of 3.1 Å per residue and the theoretical radius of gyration (R_G_) for a random coil, are given (R_G_=R_0_ N^v^ with R_0_=1.927 Å, v=0.598, and N is the number of residues). Both distances spanning the polyP regions (P11 and P10) correspond nicely to a PPII distance. The distances between residues 75 - 91, and 101 -114 are significantly shorter than a PPII helix and longer than that expected for a random coil structure.

### MD simulation of the PRD

Our previous EPR and solid-state NMR data suggested that the PRD structure in fibrils and in the soluble fusion protein is highly similar (12, 13). We therefore simulated a monomeric PRD. To compare the simulation to our solid-state NMR relaxation data recorded on HTTex1 fibrils, we fixed the N, CA, and CO atoms of the last glutamine of the polyQ domain (i.e. Q62). The C-terminus was placed in a periodic water-filled box as described in the Materials and Methods section. We then ran short initial MD simulations to test several force fields, water models, and starting structures for their ability to reproduce our DEER distances. We found that the starting structure (an extended polyproline II helix) was among the most important parameters to correctly reproduce the DEER distances. In addition, the water model was an important factor. An implicit water model and the explicit TIP3P water model with the CHARMM36 force field (44) resulted in collapsed conformations that were inconsistent with our DEER measurements. This aligns with other studies that have shown this water model is poorly suited for simulating highly extended and dynamic proteins (45). Using the AMBER ff99SB force field (35) for the protein in combination with the explicit TIP4P-D water model (34) developed specifically for intrinsically disordered proteins, led to the best fit between the DEER distances. No experimental constraints were used during these simulations, besides anchoring the N-terminus of the PRD as described above. We used this combination of force field and water model together with an extended PPII starting structure for all further simulations. We than ran two separate 800 ns simulations, the first of the HTTex1 C-terminus with a C-terminal HIS-tag and a second simulation of the C-terminus without a HIS-tag. The RMSD analysis in Figure S1 showed that both simulations had achieved equilibrium by 200 ns. All simulation frames following this timepoint were used for further analysis.

### Comparison to DEER distances

To compare the DEER distance distributions, *P*_DEER_, to the distance distributions from our MD simulations, we needed to add the corresponding MTSL labels and consider their flexibility (46). We did this by adding a set of different MTSL spin label rotamers to every 10^th^ frame (i.e. every 200 ps) of our simulations using the program RotamerConvolveMD, which is based on the MDAnalysis python package (37–39). This program also calculates a resulting distance distribution, *P*_MD_, that can be compared directly to the distance distribution *P*_DEER_. As can be seen from **Figure 3**, both the mode and the overall shape of these distance distributions are very similar.

**Figure 3:**
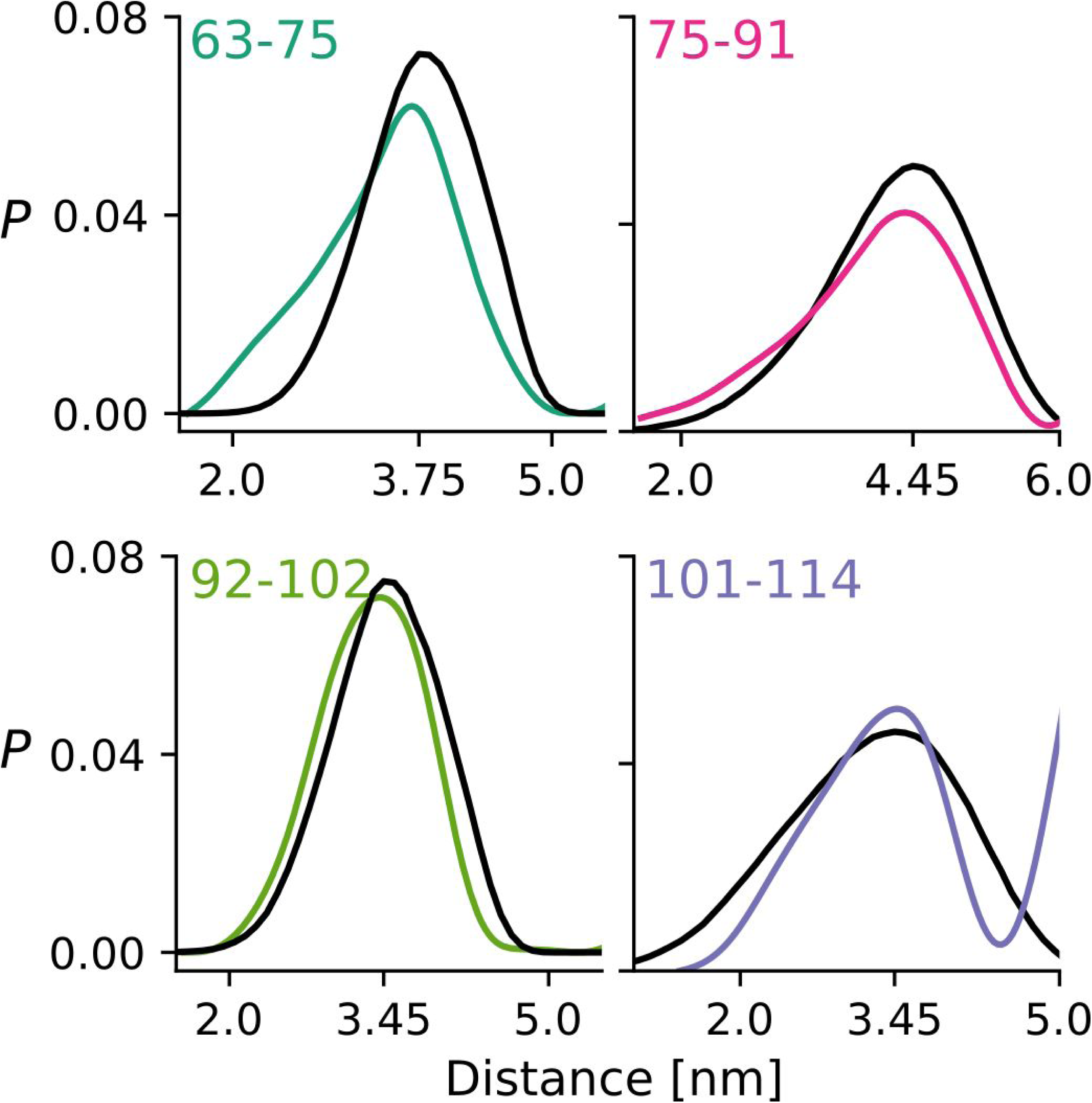
DEER distance distributions (P_DEER_) and MTSL spin label distance distributions calculated from the MD trajectory (P_MD_) correspond very well. DEER distance distributions between residue pairs as indicated are shown in colors. These correspond to distributions shown in Figure 2. Distance distributions derived from the MD simulation using the RotamerConvolveMD algorithm (P_MD_) are shown in black. The mode of P_MD_ is indicated on the x-axis.

### Comparison to NMR parameters

We recently reported the assignment of the C-terminal residues of HTTex1 fibrils starting from residue G102 (C12 region) using a combination of solution and solid-state NMR techniques. This assignment allowed us to also measure site specific ^15^N *R*_*1*_ and *R*_*2*_ relaxation rates as well as residual ^1^H-^15^N dipolar couplings that confirmed the highly dynamic nature of these residues in the context of the fibril (15). Using the traces from our MD simulations, we now calculated the *R*_*1*_ and *R*_*2*_ relaxation rates using the theoretical description by Schanda and Ernst (40) and the approach outlined in the Materials and Methods section. The comparison of the measured and theoretical relaxation rates is shown in **Figure 4**.

**Figure 4:**
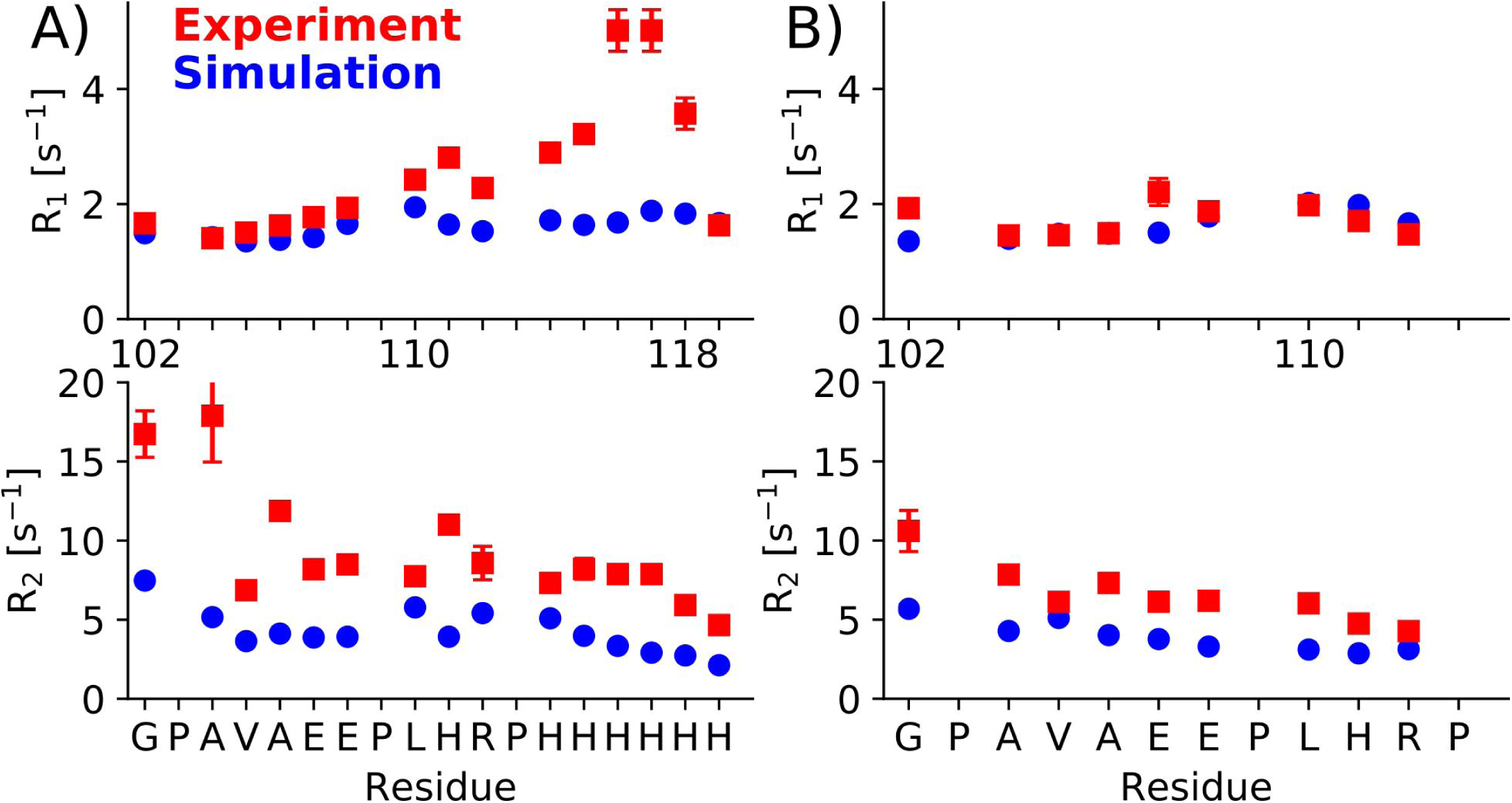
Good agreement between experimental and simulated R_1_ and R_2_ relaxation rates for the C12 region of HTTex1. A) Comparison between R_1_ and R_2_ rates that were measured from fibrils of HTTex1 with a C-terminal HIS-tag and the equivalent relaxation rates calculated from simulations of the same construct. Proline residues are omitted because of the absence of an amide proton. B) Same comparison as in A but between data and simulations of HTTex1 fibrils without a C-terminal HIS-tag. Experiment and simulations agree better for the HTTex1 fibrils without a HIS-tag presumably because of the charge repulsion of the HIS-tag in our experiments that was not captured by our simulations. In the absence of a HIS-tag, most R_1_ rates are within the margin of error of their calculated counterparts. The experimental R_2_ values are slightly higher than those calculated from simulations because of residual coherent dephasing that leads to higher apparent R_2_ rates. Experimental relaxation rates have been previously reported in Caulkins et al. 2018.

The calculated *R*_*1*_ rates for the C12 region without a HIS-tag are consistent with the experimental data and with two exceptions (G102 and E108) within the error margins of the actual rates. For the C-terminus with the HIS-tag, the calculated *R*_*1*_ rates match the experimental data at the beginning of the C12 region but show substantial differences for the residues preceding the HIS-tag and for the HIS-tag itself. These differences likely originate from the fact that the simulations were done with a non-protonated state of the HIS-tag, whereas the actually HIS-tag would have been at least partly protonated.

The calculated *R*_*2*_ rates are generally lower than the *R*_*2*_ rates measured via *R*_*1ρ*_ experiments. Again, the differences are larger for the C12 region with a HIS-tag than the C12 region without a HIS-tag, for which experimental and simulated *R*_*2*_ values correspond quite well. That the experimental *R*_*2*_ rates are larger than those calculated from the MD simulations is not surprising considering that experimental *R*_*2*_ rates usually include contributions other than transverse relaxation. In addition, we showed that there are still residual dipolar ^1^H-^15^N couplings in the C12 that are not completely averaged out by motion (15). These couplings lead to coherent dephasing and thereby an experimental overestimation of *R*_*2*_ (40).

The excellent fit between the *R*_*2*_ and *R*_*1*_ rates, especially for the C12 region without the HIS-tag, using simulations of less than 1 μs, indicates that there are no slow dynamics in this region that would not be captured by simulations that are too short. To test this interpretation, we compared *R*_*1ρ*_ relaxation rates measured at spin lock fields of 6 kHz and 18 kHz. These relaxation rates should differ (known as relaxation dispersion) if slow dynamics are present. As can be seen in ***Figure 5***, most of the *R*_*2*_ values calculated from the *R*_*1ρ*_ rates are within the error range, confirming the absence of significant slow dynamics in the C-terminus of HTTex1 fibrils.

**Figure 5:**
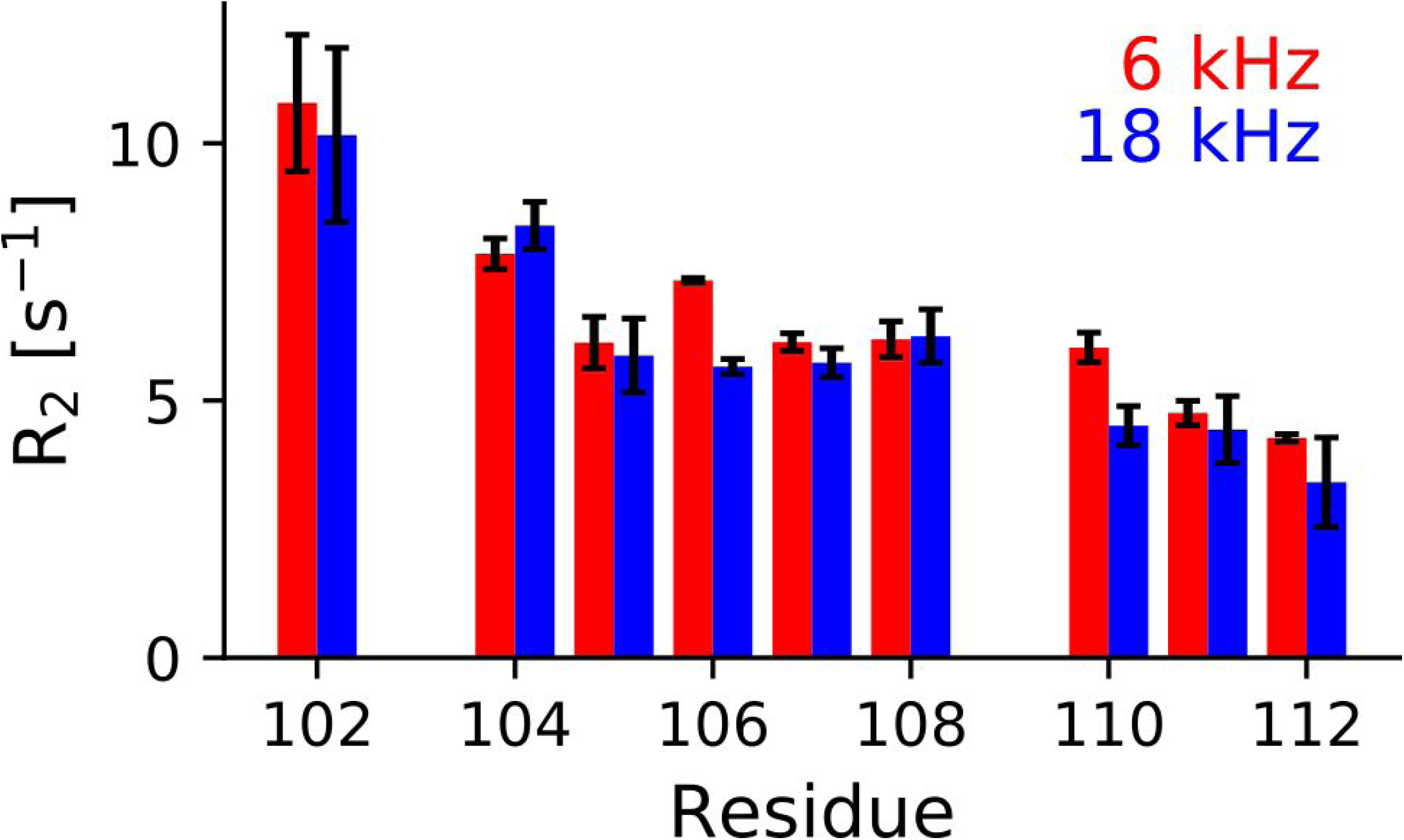
Absence of relaxation dispersion in the C12 region of HTTex1 indicates that there are no slow dynamical processes in this region. The effective R_2_ rates calculated from R_1ρ_ experiments with a spin lock rf-field strength of 6 kHz and 18 kHz are shown as red and blue bars, respectively. Except for A106 and L110 these measurements are within the margin of error each other.

The absence of slow processes in the C-terminus of HTTex1 indicates that our simulations captured the conformational space of the PRD quite well. To further confirm this, we calculated site-specific Cα chemical shifts from our simulations using the program SHIFTX2 (42). We computed the chemical shifts for each simulation frame after equilibration and calculated their average and standard deviation. The comparison of the secondary Cα shifts calculated this way, with our previously published Cα shifts (15), is shown in **Figure 6**.

**Figure 6:**
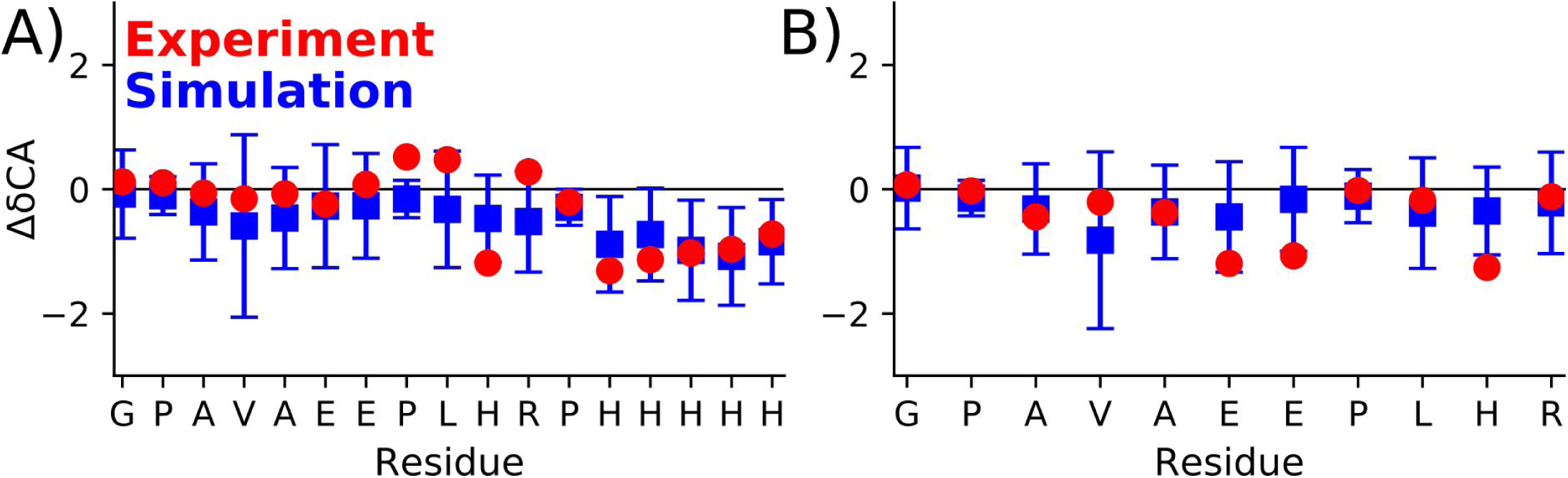
Calculated and experimental secondary Cα chemical shifts of the PRD C12 region are in good agreement. A) Average Cα secondary chemical shifts of the simulation with a HIS-tag for the C12 region are shown in blue with error bars corresponding to their standard deviation. The experimental secondary chemical shifts are shown in red. B) Average Cα secondary chemical shift of the simulation and experiments without a HIS-tag.

### Analysis of Structural Ensemble

The ability of the simulations to reproduce EPR and NMR measurements indicates that they form a representative ensemble of the structural distribution sampled by the PRD. Consequently, we analyzed the results of our simulations to gain additional insights into the structure and behavior of the PRD, with a focus on the simulation with His-tag (Figures of the simulation without the His-tag can be found in the Supporting Material).

Visual inspection of the simulations showed that the two polyP stretches remained in relatively stable PPII helices, while the L17 region connecting the two polyP stretches and the C12 region were more flexible. Because PPII helices cannot be detected by the DSSP algorithm that is based on hydrogen bond formation (47), we analyzed individual Ψ and φ angles for all residues post equilibration to confirm this observation. The average Ψ and φ angles and their standard deviation are shown in **Figure 7**. The corresponding data for the simulation without the His-tag is shown in Figure S2. All Pro residues stayed within a canonical PPII helix and showed almost no flexibility in their φ angles (48). Interestingly, Pro residues outside, or on the edges of the two polyproline stretches displayed more Ψ angle flexibility. Almost all non-Pro residues adopted dihedral angles that were between the canonical angles for a β-sheet and a PPII helix. These angles are consistent with our observation that the C-terminus remains in a relatively extended conformation. Two exceptions to these extended dihedral angles were A83 and L86 in the no-His-tag simulation, which had average dihedral angles between those found in a β-sheet and an α-helix. The difference in the dihedral angles of A83 and L86 is one of the few major differences between the two simulations.

**Figure 7:**
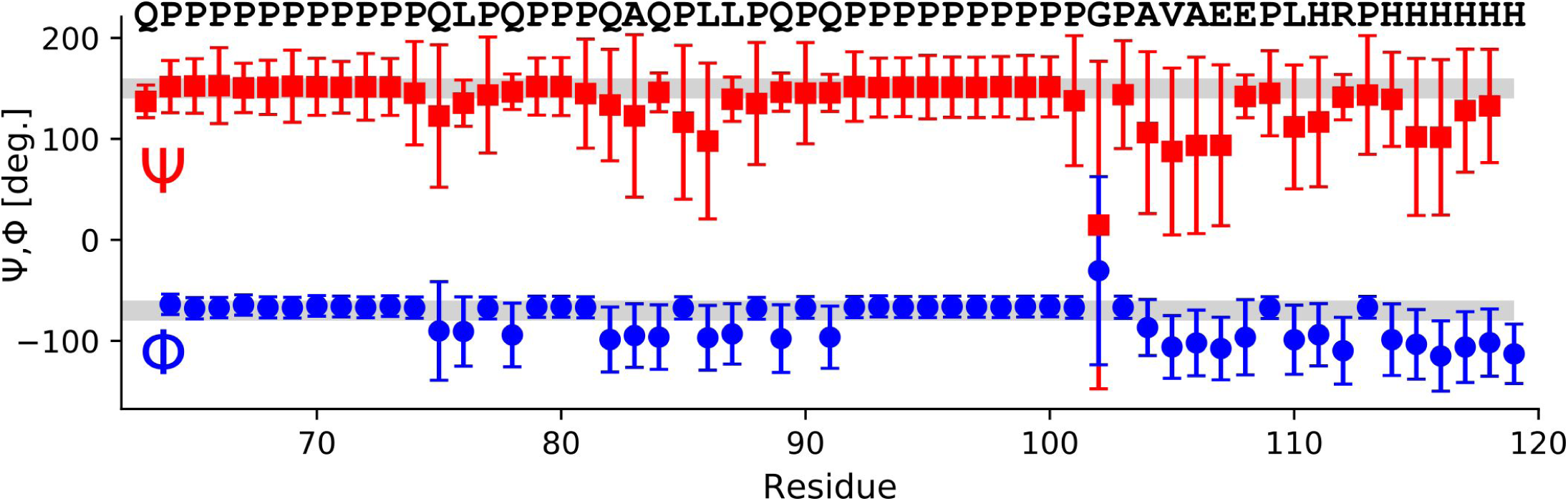
The variation in φ and Ψ angles demonstrates relatively stable PPII helices for the polyP stretches and an increase in disorder for the L17 and C12 region. For the simulation with a HIS-tag, average per residue φ and Ψ angles with their standard deviations are shown in red and blue respectively. The φ and Ψ angles for an idealized PPII helix are indicated with gray bars. The very large error bars for residue G102 shows that the C12 region can rotate relatively freely around this residue.

Another important exception from generally extended dihedral angles is G102 showing significant flexibility in its dihedral angles as illustrated by the large error bars in **Figure 7**. This is not surprising given Gly’s nature, but it is worth noting that all residues following G102 show much higher degrees of variation and disorder than the residues preceding G102.

But how flexible is the L17 region, does it allow the PRD to fold back onto itself? In order to address this question, we plotted Cα-Cα distances over the course of the simulation with His-tag. As can be seen from **Figure 8**, the distance over the N17 region (75-91) is compatible with an extended PPII helix for most of the simulation with clear exceptions in which this region kinks, shortening its overall extension. In contrast, the two PPII stretches (63-75 and 91-102) remain essentially fixed in an extended PPII helical conformation. The C12 region (101-114) is the most flexible part of the PRD. Although it stays relatively extended throughout the simulation, it is generally not in an extended, PPII conformation. The overall extension of the PRD, represented by the distance between Q63 to G102, is often correlated with the bending of the L17 region (see gray boxes in **Figure 8**).

**Figure 8:**
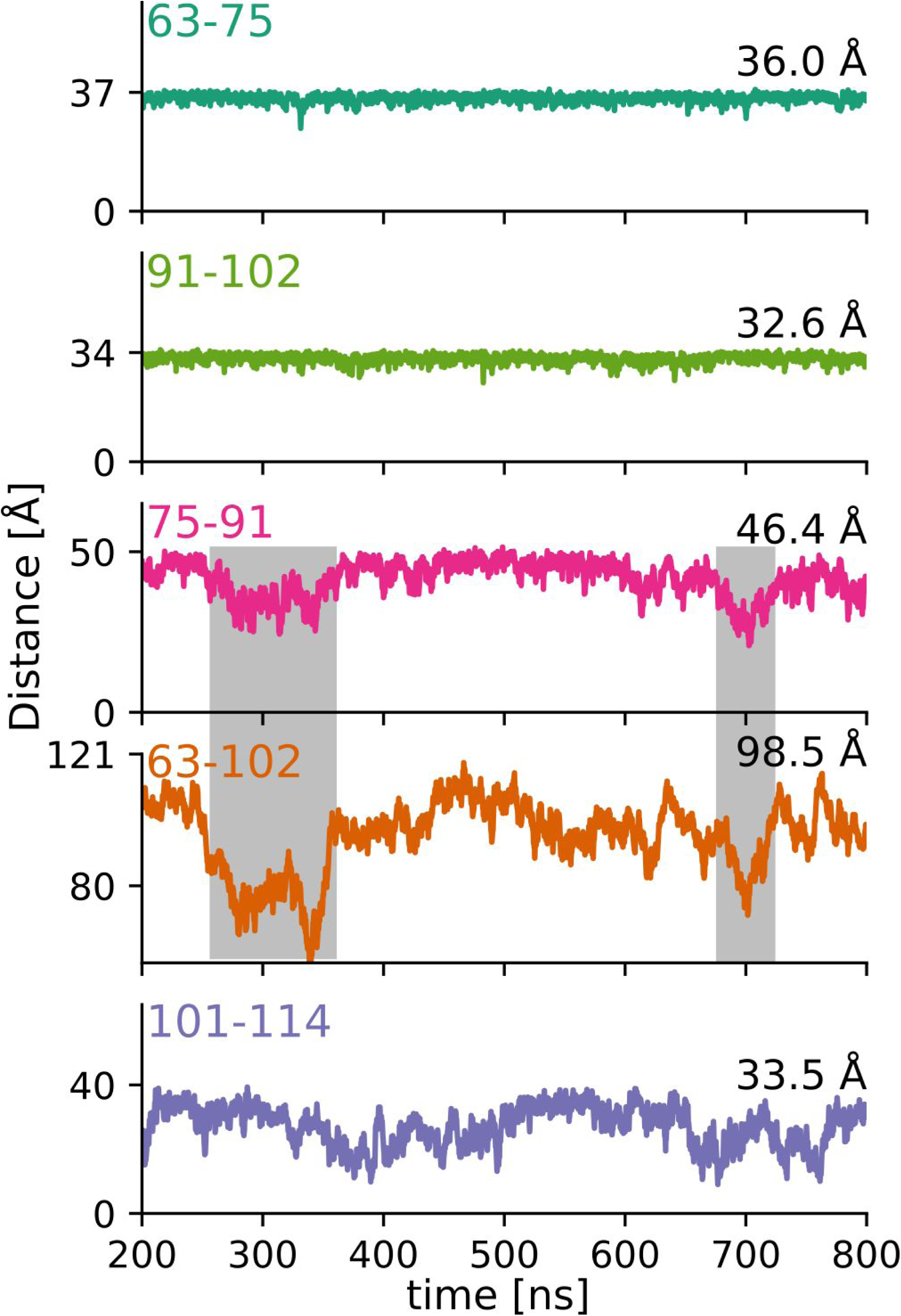
The L17 region is responsible for the overall kinks of the PRD. Change in Cα-Cα distances (as defined in **Figure 1**) over time. The upper y-axis tick indicates an ideal polyproline II helical distance (see **Figure 2f**), the mode of simulated Cα-Cα distance distribution is indicated on the right. All distance variations are plotted with comparable y-axis scaling. Both polyP stretches stay in an extended PPII conformation for most of the simulation. The L17 region is a less ideal PPII helix. Its deviations from an extended PPII conformation induce an overall kink in the PRD as illustrated by the correlated shortening of distance 63-102 (gray boxes). The C12 region is the most flexible and although still relatively extended, deviates from a PPII helix most of the time.

To get a better sense of the conformational space occupied by the PRD, we clustered the MD trajectory using the K-means algorithm. We determined a suitable number of clusters by dividing the trajectory into 2 to 20 clusters and calculated the pSF value and SSR/SST ratio for each of these divisions (see Figure S3). At a suitable number of clusters, pSF reaches a local maximum and the SSR/SST ratio starts to plateau (49). In our case this was the case at 3 clusters. The centroids of each of these 3 clusters together with a schematic of the PRD are shown in **Figure 9**. All centroids have extended P11 and P10 regions, and a relatively disordered C12 region in common. The conformation of the L17 region determines the overall shape of the domain. Consequently, the PRD is relatively extended in centroid 2 where the L17 domain is extended as well, less extended in centroid 3 where the L17 regions adopts an s-shaped conformation, and significantly shorted in centroid 1 where the L17 is kinked. This analysis further confirms that although the PRD of HTTex1 is predominantly in a PPII helical conformation, it has the ability to kink at the L17 region.

**Figure 9:**
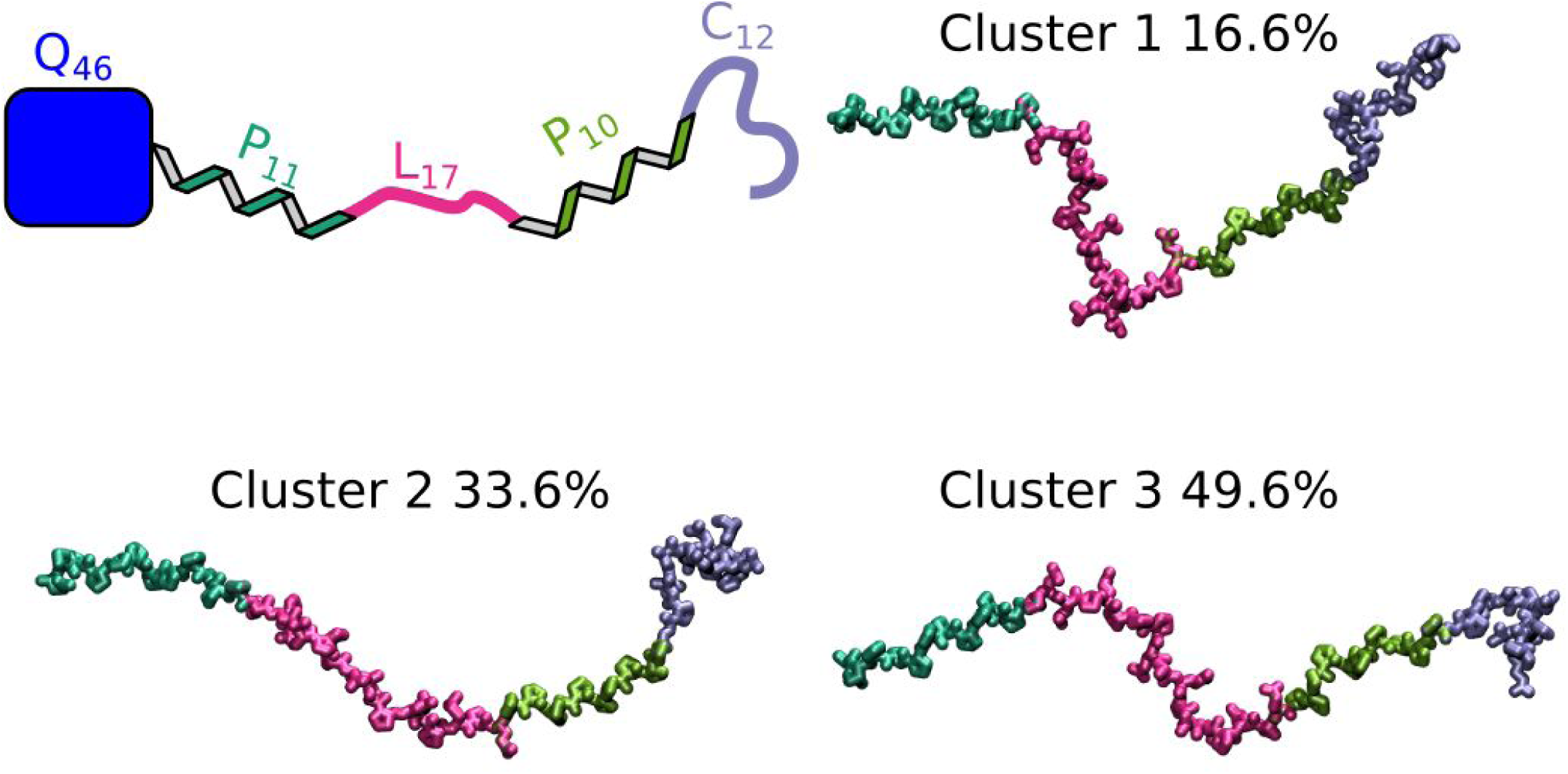
Cluster analysis of MD trajectory showing how the conformation of the L17 region defines the overall structure of the PRD. A) Schematic of the PRD connected to the polyQ fibril core of HTTex1 fibrils. B) Centroids of the K-means cluster analysis. The percentages of total frames they represent are indicated. Clusters 2 and 3 are relatively extended, whereas cluster 1, having a strong kink in the L17 region, is less extended.

## Discussion

The present study showed that MD simulations using the AMBER ff99SB force field with the TIP4P-D water model led to trajectories for the PRD of HTTex1 that correlates very well with EPR DEER distance distributions, NMR ^15^N relaxation rates, and NMR Cα chemical shifts. Overall, the PRD stays relatively extended throughout the simulations with two stable PPII helices, P10 and P11, a more variable L17 linker region, and a very flexible C-terminal C12 region. Nevertheless, the L17 and to a lesser extend the C12 region have average dihedral angles compatible with a PPII helical or β-sheet conformation and are extended for most of the simulation, indicating that these regions are rather imperfect PPII helices than completely disordered.

The large distribution of dihedral angles for G102 indicates that this residue may have a role in separating the Pro-rich area from subsequent HTT domains and effectively terminates the order imposed by the Pro residues. In addition, the flexibility of G102 explains why the C12 region could be detected in our HSQC spectra in the absence of perdeuteration and at relatively slow MAS frequencies as reported previously (15). G102 allows the residues in the C12 region to rotate relatively freely, resulting in an order parameter that is essentially zero and an almost complete averaging of the ^1^H-^15^N dipolar couplings that allowed the direct ^1^H detection in our NMR experiments. In contrast, the preceding polyP regions could not be detected in the ^1^H-^15^N HSQC experiment because of the absence of an amide proton in the polyP stretches and the reduced flexibility of the L17 region that was not enough to average the H-N dipolar coupling such that ^1^H detected experiments became feasible under the conditions used. It is interesting to note that the C12 region is evolutionary well conserved and we speculate that it might serve as a dynamic linker to the well-structured and conserved first HEAT repeat of HTT (50, 51).

Our finding that the PRD is mostly extended is compatible with the tadpole model of the HTTex1 monomer by Newcombe and co-workers in which the N17 and polyQ domains are more compact and the PRD forms the extended tail of a tadpole-like structure (24). In contrast to our simulations, their modeling approach was based on Monte Carlo simulations and an implicit water model optimized for intrinsically disordered proteins as implemented in the ABSINTH program (52). This, and the fact that they assumed the His residues in the C12 region to be protonated, likely explains some of the differences with our results. Namely, the propensity of the C12 region to form an α-helix in their simulation.

For the two polyP regions, the mode of the Cα-Cα distance distribution of our simulation (see **Figure 8**) is shorter than an idealized PPII helix but also a bit longer than what was described by Radhakrishnan and co-workers using the ABSINTH algorithm (53(53)). This mode increases after addition of spin labels using the RotamerConvolveMD algorithm because for both distances (63-75, 91-102) the labels point into different directions relative to the helix norm. The relatively good fit to the EPR distance distributions suggests that our simulations created a valuable model of the two polyP regions.

Our simulations focus on a single PRD, because our previous EPR and NMR data showed that the structure of this domain is very similar in the soluble fusion protein and HTTex1 fibrils (12, 13). That our simulations reproduce the solid-state NMR data from the C12 region in HTTex1 fibrils further supports this finding, suggesting that this region is not affected by potential PRD-PRD interactions inside the fibril. This seems to be also true for the rest of the PRD. The DEER distances within the PRD of the fusion protein and different fibril types, which we reported previously (25), are very similar.

Our results are consistent with the ability of the PRD to inhibit fibril formation. Because the PRD is dynamic in both soluble HTTex1 and the fibril, it likely counteracts fibril formation by imposing a PPII conformation on the polyQ domain rather than creating an entropic penalty from being placed into the fibril (16, 54).

Many HTTex1 specific antibodies bind the PRD. MW7 and 4C9 bind the polyP regions, MW8 binds the C12 region, and PHP1 and PHP2 bind the L17 region (55–57). Interestingly, all of these antibodies are fibril specific and only weakly bind to soluble HTTex1. The present work shows that the polyP, L17, and C12 regions not only differ in sequence but also in their degree of dynamics and deviation from a PPII structural motif. Therefore, it is possible that these epitopes are not only distinguished by their amino acid sequence but also by their structural preference. Similarly, we hope that the PRD model presented in this paper will help understand how some fibril-specific HTTex1 interactors such as chaperones (58) bind.

## Conclusions

We simulated the PRD of HTTex1 in fibrils using the AMBER ff99SB force field and TIP4P-D water model. These simulations accurately predicted our EPR and solid-state NMR data indicating that the PRD does not undergo slow processes that would not be captured by less than1 μs of simulation. The PRD adopted a predominantly PPII helical conformation for most of the MD trajectory. The two polyP regions formed stable PPII helices, the L17 region formed an imperfect PPII helix, and the C12 region only loosely maintained the PPII helical conformation. G102, at the beginning of the C12 region, was the most flexible residue, separating the PRD from the following highly conserved regions of HTT. Besides these structural insights, our study shows that modern MD methods in combination with EPR and solid-state NMR can accurately characterize intrinsic disorder in non-soluble proteins.

## Author Contributions

A.S.F. ran and analyzed MD simulations and co-wrote the paper. J.M.B. made EPR samples. J.V. measured EPR data. S.P. performed the cluster analysis of the MD data. R.L. coordinated the EPR work and its interpretation. A.B.S. conceived the study, recorded the NMR data, analyzed MD simulations, and co-wrote the manuscript.

## Acknowledgments

A.B.S and R.L. would like to acknowledge funding from the National Institutes of Health (R01NS084345, R01GM110521), and the CHDI Foundation (Award A-12640). A.S.F would like to acknowledge funding from the National Institutes of Health (F31GM120858). J.M.B. would like to acknowledge a USC-CONACYT fellowship.

